# Distinguishing the roles of dorsolateral and anterior PFC in visual metacognition

**DOI:** 10.1101/280081

**Authors:** Medha Shekhar, Dobromir Rahnev

**Affiliations:** School of Psychology, Georgia Institute of Technology, Atlanta, GA

## Abstract

Visual metacognition depends on regions within the prefrontal cortex. Two areas in particular have been repeatedly implicated: the dorsolateral prefrontal cortex (DLPFC) and the anterior prefrontal cortex (aPFC). However, it is still unclear what the function of each of these areas is and how they differ from each other. To establish the specific roles of DLPFC and aPFC in metacognition, we employed online transcranial magnetic stimulation (TMS) to causally interfere with their functioning during confidence generation. Human subjects from both sexes performed a perceptual decision-making task and provided confidence ratings. We found a clear dissociation between the two areas: DLPFC TMS lowered confidence ratings, whereas aPFC TMS increased metacognitive ability but only for the second half of the experimental blocks. These results support a functional architecture where DLPFC reads out the strength of the sensory evidence and relays it to aPFC, which makes the confidence judgement by potentially incorporating additional, non-perceptual information. Indeed, simulations from a model that incorporates these putative DLPFC and aPFC functions reproduced our behavioral results. These findings establish DLPFC and aPFC as distinct nodes in a metacognitive network and suggest specific contributions from each of these regions to confidence generation.

**Significance**

The prefrontal cortex (PFC) is known to be critical for metacognition. Two of its sub-regions - dorsolateral PFC (DLPFC) and anterior PFC (aPFC) - have specifically been implicated in confidence generation. However, it is unclear if these regions have distinct functions related to the underlying metacognitive computation. Using a causal intervention with transcranial magnetic stimulation (TMS), we demonstrate that DLPFC and aPFC have dissociable contributions: targeting DLPFC decreased average confidence ratings, while targeting aPFC specifically affected metacognitive scores. Based on these results, we postulated specific functions for DLPFC and aPFC in metacognitive computation and corroborated them using a computational model that reproduced our results. Our causal results reveal the existence of a specialized modular organization in PFC for confidence generation.

## Introduction

Metacognition, or the ability to assess the quality of our decisions, is crucial for effective decision making (Metcalfe and Shimamura, 1994; Koriat, 2007). However, despite the critical influence of metacognition on our actions and decisions (Nelson and Narens, 1990; Shimamura, 2000a; Koriat, 2007; Fleming et al., 2012a; Yeung and Summerfield, 2012), its neural bases are still not fully elucidated (Shimamura, 2000a; Fleming et al., 2012b). Early studies pointed to a central role of the prefrontal cortex (PFC) based on findings of impaired metacognition in patients with damage to the frontal lobe (Shimamura and Squire, 1986; Janowsky et al., 1989; Shimamura, 2000a). More recent research has implicated two specific PFC sub-regions - the dorsolateral prefrontal cortex (DLPFC) and the anterior prefrontal cortex (aPFC) (Fleming & Dolan, 2012).

Activity in DLPFC has been linked to the level of reported confidence. Indeed, studies employing functional magnetic resonance imaging (fMRI) have consistently shown that activity within DLPFC tracks confidence levels during metacognitive computations (Fleck, et al., 2005; Henson et al., 2000; Lau & Passingham, 2006; Morales,et al., 2017).

On the other hand, aPFC has been specifically linked to subjects′ metacognitive ability. For example, structural imaging studies have found that grey matter volume in aPFC correlates with individual metacognitive ability (Fleming et al., 2010; Yokoyama et al., 2010; McCurdy et al., 2013; Allen et al., 2017). Similarly, studies using fMRI show that aPFC activity is modulated by the reliability of confidence judgments (Yokoyama et al., 2010; Fleming et al., 2012b; Morales et al., 2017). Finally, metacognitive scores are affected by both lesions (Fleming et al., 2014) and transcranial magnetic stimulation (Rahnev et al., 2016; Ryals et al., 2016) to aPFC.

Based on the findings above, we hypothesized specific functions for DLPFC and aPFC in confidence computation. We propose that DLPFC reads out the strength of the sensory signal and relays it to aPFC. The readout of the sensory signal determined by DLPFC conveys how much information was available for the sensory decision. aPFC subsequently integrates this readout with additional, non-perceptual factors and translates all this information into a confidence judgment **(Figure 1).** Disrupted readout of the sensory signal by DLPFC would, on average, convey that less information was available for the sensory decision. In turn, aPFC would translate such disrupted readout into lower confidence ratings. This architecture is consistent with prior findings since reading out the sensory signal strength would link DLPFC activity with confidence level, while making the confidence judgment would link aPFC activity with metacognitive ability. In addition, (Fleming et al., 2012b) observed that connectivity between aPFC and DLPFC increased during metacognitive reports, suggesting active communication between the two regions during confidence computation.

**Figure 1:**
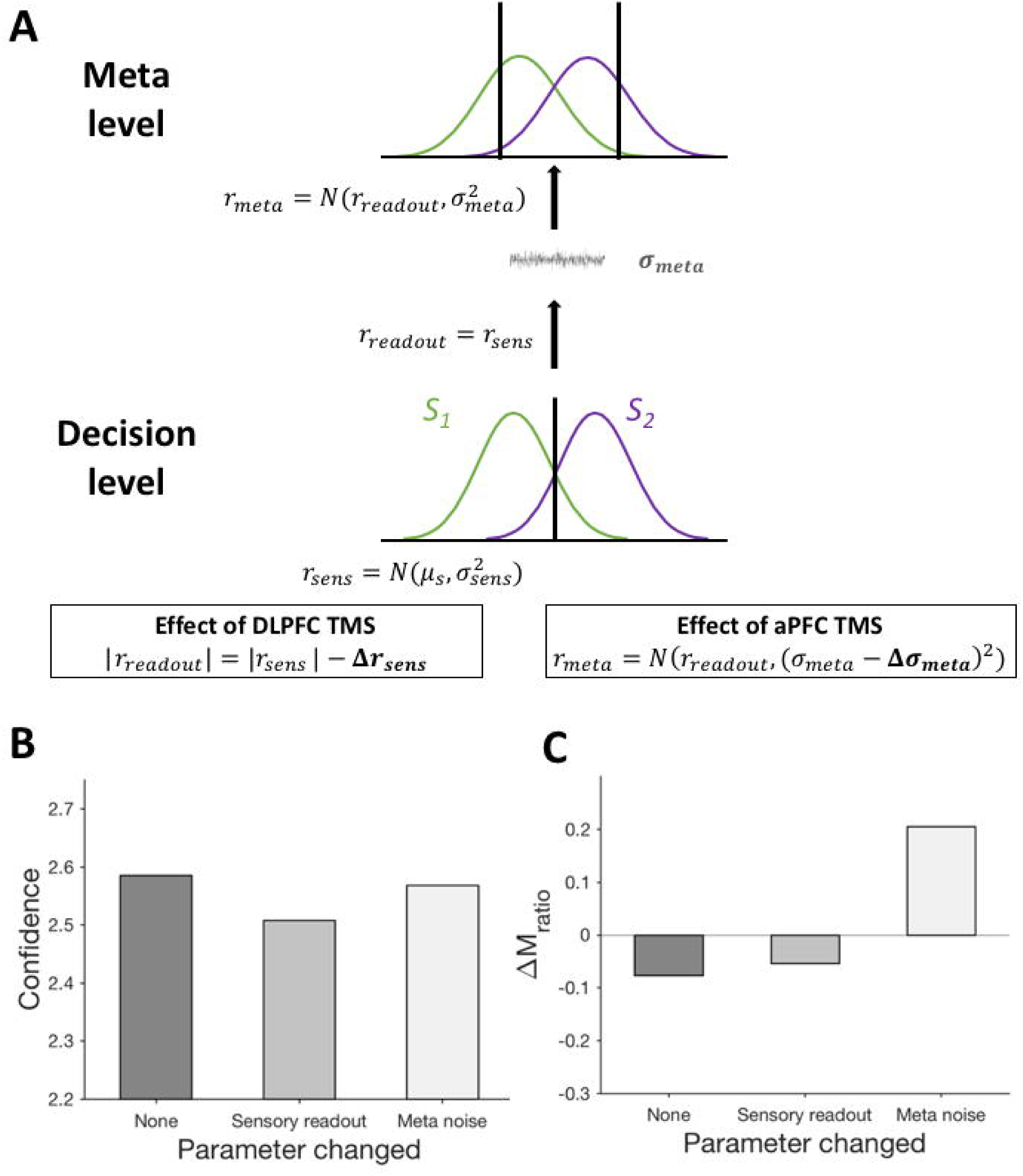
Hypothetical neural mechanism of confidence computation. Based on prior literature, we postulated the following neural mechanism for the roles of DLPFC and aPFC in confidence computation. DLPFC reads out the strength of the sensory signal and relays it to aPFC. On the other hand, aPFC translates this readout into a confidence judgment after incorporating additional, non-perceptual factors. The strength of the sensory signal that is read out by DLPFC on a particular trial is related to the level of confidence on that trial. On average, disrupting the readout would convey that less evidence was available for the perceptual decision compared to the evidence that was actually available. Such disrupted readout would be translated by aPFC into a lower confidence rating. Therefore, impaired DLPFC functioning leading to poor quality readouts would convey that the sensory information was more ambiguous than it really is and would result in lower confidence ratings. In contrast, impaired aPFC functioning would alter how aPFC transforms the sensory readout relayed from DLPFC into a confidence judgement and, therefore, alter metacognitive performance.

We tested our hypothesis regarding the putative functions of DLPFC and aPFC in confidence computation by employing online transcranial magnetic stimulation (TMS). Previous TMS studies on visual metacognition (Rahnev et al., 2016; Rounis et al., 2010; Ryals et al., 2016) used offline approaches that inhibit activity for an extended period of time. These studies showed little or no modulation of overall confidence level presumably because subjects had time to re-calibrate their confidence judgments. To address this issue, we applied online TMS in short blocks to avoid behavioral compensation.

Based on our hypothesis about the functions of DLPFC and aPFC, we predicted that TMS to DLPFC would affect subjects′ overall confidence level, while TMS to aPFC would affect metacognitive ability. The results confirmed these predictions: TMS to DLPFC decreased confidence, whereas TMS to aPFC increased metacognitive ability but only for the second half of blocks. Further, we confirmed that these results can be reproduced by a model in which TMS to DLPFC affected the readout of the sensory information, while aPFC TMS affected the noise within the metacognitive computation itself. Our findings demonstrate that DLPFC and aPFC have distinct functions in visual metacognition and suggest a specific mechanistic role for each.

## Methods

### Subjects

A total of 21 subjects completed the study (13 females and 8 males, average age = 22 years, age range = 18-32 years). Three subjects were excluded from analyses. For one subject, the sensors registering the subject′s brain to their MRI shifted mid-session, which likely resulted in imprecise TMS target localization. The other two subjects were excluded due to poor performance or excessive number of interruptions due to discomfort. All subjects were right handed and had normal or corrected-to-normal vision.

### Session sequence

We collected data for our experiment over two sessions, which were held on separate days. By dividing data collection into two days, we were able to collect more data while keeping the session short enough to avoid fatigue.

Day 1 started with a short training on the behavioral task, followed by a staircasing procedure used to identify the contrast of the stimulus to be used for the main experiment. After subjects completed the staircasing, we determined the amplitude of TMS stimulation to use and started the main experiment.

The main experiment consisted of four runs of three blocks each. For each of the three blocks within each run, we stimulated one of three regions - dorsolateral prefrontal cortex (DLPFC), anterior prefrontal cortex (aPFC), or the somatosensory cortex (S1; which served as the control site) - in a pseudo-random order such that all the three sites were stimulated once within each run. The first run was a practice run and was shorter than the others. It was included to accustom subjects to receiving TMS to the different brain regions and minimize chances of the TMS pulse evoking a startle response during the main trials. The blocks from the practice run consisted of 16 trials each and were excluded from further analyses. All other blocks consisted of 40 trials each. Therefore, subjects completed a total of 408 trials during each session.

During Day 1, subjects underwent a behavioral training procedure without TMS. The training session started with high stimulus contrast values (40%) and gradually presented lower contrast values (the last block included contrast values of 4%). Subjects were given trial-by-trial feedback on their performance during this training period.

At the end of the training, subjects completed a 3-down-1-up staircasing procedure consisting of trials without feedback. The 3-down-1-up procedure is a variant of the up-down transformed response method used for adaptive estimation of stimulus thresholds (Macmillan and Creelman 2005). This procedure yielded a contrast value for the stimulus (mean = 6.64% and SD = 0.96%) that was expected to result in an accuracy of 79% (Macmillan and Creelman 2005; observed mean = 79.6% and SD = 5.8%). We used the contrast value obtained from this procedure for the rest of the experiment.

Day 2 was identical to Day 1 except that subjects did not have to undergo behavioral training or staircasing (we used the same stimulus contrast as in Day 1).

### Task

Each trial began with subjects fixating on a small white dot (size = 0.05°) at the center of the screen for 500 ms followed by presentation of the stimulus for 100 ms. The stimulus was a Gabor patch (diameter = 3°) oriented either to the right (clockwise, 45°) or to the left (counterclockwise, 135°) of the vertical and was superimposed on a noisy background. Subjects′ task was to determine the orientation of the Gabor patch while simultaneously rating their confidence on a 4-point scale (where 1 corresponds to the lowest confidence rating and 4 corresponds to the highest confidence rating) via a single key press. Subjects placed their fingers of each hand on a standard keyboard. The four fingers of the left hand were mapped to the four confidence responses for the left-tilted stimulus, while the four fingers of the right hand were mapped to the four confidence responses for the right-tilted stimulus. For each hand, the index finger indicated a confidence of 1, while the little (pinky) finger indicated a confidence of 4. The orientation of the stimulus (left/right) was chosen randomly on each trial.

We delivered TMS on each trial as a train of three pulses delivered 250, 350, and 450 ms after stimulus onset. We chose this timing so that it coincides with the presumed time window of confidence computation. While there is no clear data on the precise time window when confidence is computed, neuronal recordings from monkeys suggest that the discrimination response emerges ∼200 ms following stimulus onset (Siegel et al., 2015), placing confidence computation in human PFC no earlier than 200 ms. To estimate the length of the time window, we collected pilot data from an identical discrimination task where subjects made two responses: their first response indicated the tilt (left/right) of the Gabor patch and their second response indicated their confidence level on a 4-point scale. Analysis of these data showed that subjects typically take ∼500 ms to give their confidence response, following the discrimination response. After roughly accounting for motor preparation (∼200 ms), we estimated that the actual duration of confidence computation is about 300 ms. Based on this estimation, we timed our TMS pulses so that they targeted a time window that started 250 ms following stimulus onset and spanned the next 200 ms.

### Apparatus

Stimuli were generated using Psychophysics Toolbox in MATLAB (MathWorks). During the training and the main experiment, subjects were seated in a dim room and were positioned 60 cm away from the computer screen (21.5-inch display, 1920 × 1080 pixel resolution, 60 Hz refresh rate).

### Defining regions of interest (ROIs) for TMS targeting

We defined three sites as targets for TMS: dorsolateral prefrontal cortex (DLPFC), anterior prefrontal cortex (aPFC) and the somatosensory cortex (S1; control site). Based on previous studies (Fleming et al., 2010, 2012b; Yokoyama et al., 2010; Rahnev et al., 2016), aPFC was localized at [28, 56, 26]. We localized DLPFC immediately posterior to aPFC (at a distance of 2.6 cm posterior to aPFC) and used [28, 30, 38] as the target coordinates. For S1, we used [20, −39, 70] as the putative coordinates (Rahnev et al., 2016) but the actual location of stimulation was adjusted based on Si′s known anatomical location in the postcentral gyrus. As in previous work (Rahnev et al., 2016), all regions were defined in the right hemisphere because the right hemisphere is dominant for visual processing (Hellige, 1996).

We defined the ROIs on the anatomical MRI scans of each subject. These scans were obtained during previous studies conducted in the lab. In order to determine the subject-specific location for stimulation, we back-normalized the coordinates above to the subject′s native space. We created ROIs as 5-mm spheres and their centers were set as targets to guide the placement of the TMS coil. In some cases, the ROIs produced via back-normalization appeared shifted with respect to the expected anatomical location. In such cases, we switched to an alternate method of defining ROI locations. The neural navigator software, *TMSNavigator* (Localite), contains a built-in program for defining a Talairach coordinate system on a subject′s MRI that is based on the location of the anterior commissure, the posterior commissure, and the vertex. After these structures are manually identified on an MRI scan, the software generates a Talairach grid, which can be adjusted so that it encloses the whole brain. This grid allows transformation of coordinates between the subject′s native coordinate space and the MNI coordinate space.

### TMS setup

TMS was delivered with a magnetic stimulator (MagPro R100, MagVenture), using a figure of eight coil with a diameter of 75 mm.

We determined the resting motor threshold (RMT), immediately prior to starting the main experiment. In order to localize the motor cortex, we marked its putative location and applied supra-threshold single pulses around that location. We determined the location of the right motor cortex as the region that induced maximal twitches of the fingers in the left hand. Then, using this location as the target, we determined the RMT using an adaptive parameter estimation by sequential testing (PEST) procedure (Borckardt et al., 2006). For three subjects, we were unable to reliably estimate RMT, even at amplitudes as high as 80. Therefore, for these subjects we chose to determine the active motor threshold (AMT) instead, which is lower than RMT and could be found reliably. Motor thresholding was done separately for both days (average for Day 1 = 59.94, average for Day 2 = 58.28), to control for non-specific factors, which can influence the TMS response (Ridding and Ziemann, 2010).

The TMS coil was positioned on the previously-defined ROIs using a neural navigation system (TMS Navigator, Localité). The coil was oriented tangential to the skull and in such a way that the magnetic field induced was orthogonal to the skull. Stimulation was delivered at 90% of the individual resting motor threshold (RMT). In some cases when the stimulation intensity was uncomfortable to the subject, it was reduced to ∼85% (2 subjects) or ∼80% (3 subjects) of RMT depending on the individual′s comfort level. No arm or leg movements were elicited by stimulation of any of the three sites.

### Analyses

We analyzed the data for two separate measures: average confidence and metacognitive ability. To compute the average confidence, we simply calculated the average of all confidence ratings within each TMS condition. We quantified metacognitive ability using the measure *M*_*ratio*_ developed by Maniscalco & Lau (2012). *M* _*ratio*_ is derived from signal detection theoretical modeling of the observer′s decision and confidence responses. It is the ratio of two measures - the observer′s metacognitive sensitivity (*meta-d′ -* ability to discriminate between correct and incorrect responses) and the observer′s stimulus sensitivity (*d′ -* ability to discriminate between the two stimulus classes). The ratio of *meta-d′* to *d′* factors out the contribution of stimulus sensitivity towards metacognitive performance and captures the efficiency of the observer′s metacognitive processes (Fleming and Lau, 2014).

We compared the effect of TMS on confidence and metacognitive ability between the three TMS conditions (DLPFC, aPFC and S1) using one-way repeated measures ANOVAs. Additionally, we analyzed the interaction between time (within a block) and TMS location by splitting each block into first (trials 1-20) and second (trials 21-40) halves and performing a 2-way repeated measures ANOVA. Direct comparisons between regions were made using paired t-tests.

Splitting blocks in halves and analyzing each half separately may decrease the stability of the *M*_*ratio*_ estimates. To confirm that our *M*_*ratio*_ estimates were not unacceptably variable, we tested whether splitting blocks in half had a significant influence on the variance of *M*_*ratio*_ scores. We verified that all groups of A/W,_0_ values (coming from first half, second half and the whole block) were normally distributed and used the F-test of equality of variance to test whether the two distributions came from populations with different variances. First, we compared the population variance of *M*_*ratio*_ scores between the first and the second halves (after pooling /VW,o scores obtained from all three TMS conditions). The F-test showed that the between-subject variance of A/W,_0_ was not significantly different between the two halves of the blocks (F = 0.72, P = 0.23). Next, we pooled the *M*_*ratio*_ scores from both halves and compared their variance against *M*_*ratio*_ scores obtained from combining trials from both the halves. The F-test revealed no significant difference between the variance of these two populations too (F = 0.71, P = 0.16). In addition, we confirmed that the number of zero-cell counts in the accuracy/confidence matrix (that is, the number of confidence-accuracy combinations - such as incorrect trials with confidence of 4 - that never appeared) were similar between the two halves for all three TMS conditions. When such zero-cell counts did occur, we applied the same default correction for such zero-cell counts (Maniscalco & Lau, 2012) uniformly across all the conditions.

### General model architecture

Our results showed that TMS to each prefrontal site affected one specific aspect of confidence ratings - either their average value or their reliability in predicting accuracy. Our neural mechanism implies that the change in average confidence was due to TMS affecting the readout of the sensory signal and the change in metacognitive ability was caused by TMS affecting the efficiency of the metacognitive evaluation.

To assess our proposed neural mechanism, we performed simulations of a confidence generation model that incorporated our hypothesized TMS effects. It should be noted that we could not use previous approaches such as the existing procedure for estimating metacognitive sensitivity *(meta-d′),* which is built on a signal detection theoretical (SDT) framework for modeling perceptual decisions (Maniscalco and Lau, 2012). The reason is that although this procedure allows for the estimation of metacognitive performance, it is not a generative model and does not specify how the confidence data actually come about. On the other hand, to simulate confidence data, we needed a generative model. We sought to build a generative model that preserves the assumptions of SDT (Green and Swets, 1966) at the level of the perceptual decisions but also allows us to explicitly model the transformations to the sensory signal that are responsible for generating the confidence ratings. The simplest way to model the transformation of the sensory signal at the metacognitive level is to postulate the existence of metacognitive noise that corrupts the sensory signal as done previously by the creators of the *meta-d′* measure (Maniscalco and Lau, 2014), us (Rahnev et al., 2016; Bang et al., 2017) and others (Mueller and Weideman, 2008; Jang et al., 2012; De Martino et al., 2013; van den Berg et al., 2017).

Our generative model assumes that perceptual decisions and confidence ratings are the result of a hierarchical process consisting of two levels: an object level, which generates the discrimination response, and a meta level, which generates the confidence response. At the object level, the presented stimulus produces a sensory response corrupted by Gaussian noise. We modeled the two Gaussian distributions arising from the two stimulus classes (left/right tilted Gabor patches) such that the left-tilted stimuli produce a sensory response 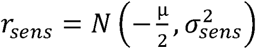 and the right-tilted stimuli produce a sensory response *r*_*sens* =_ 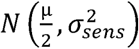 Note that the distance between these distributions is μ and the stimulus sensitivity can be expressed as: 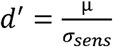. A copy of this sensory response, *r*_*sens*_, gets transferred to the meta level as a readout of the sensory signal strength, *r*_*readout*_, where it is further corrupted by metacognitive Gaussian noise such that the metacognitive response is given by the formula 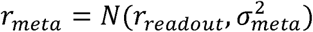.

To simulate how subjects make perceptual and confidence responses on each trial, we specified a decision criterion, c_0_, and confidence criteria, c_-n_, c__n+1_,…, *c*_*sub>t*_*, c*_t_,…, *c*_*n*_*_*_*t*_*, c*_*n*_, where n is number of ratings on the confidence scale (in our case, n = 4). The criteria *c*_*t*_ were monotonically increasing with c_-n_ = —∞ and *c*_*n*_ *=* ∞.

The object-level decisions were made based on a comparison of *r*_*sens*_ with c_0_. For trials in which *r*_*sens*_ > c_0_, the response was given as “right”; otherwise, the response given was “left. ” Confidence responses were based on *r*_*meta*_ such that values falling within the interval [_*Ci*_, c_i+1_) resulted in a confidence of *i* + 1, when *i* ≥ 0 and confidence of - *i,* when *i ≤* —1, where *i* takes integer values ranging from −4 to 3. In cases in which *r*_*sens*_ and *r*_*meta*_ fell on different sides of the decision criterion c_0_, we constrained the confidence response to equal 1.

Finally, our data showed the existence of small (and non-significant) decrease in *M*_*ratio*_ for the second half of blocks in the S1 and DLPFC TMS conditions. This effect parallels recent findings that metacognitive ability may decrease in second half of blocks due to fatigue (Maniscalco et al., 2017). To model this effect, we allowed a_me_ta to increase in the second half of all blocks by a value controlled by the parameter *Δ σ*_*meta*_*_* _*base*_

Our computational model can be related to our hypothesized neural mechanism about the roles of DLPFC and aPFC in confidence computation. According to the neural mechanism that we proposed, the sensory signal strength is read out by DLPFC. Here, we model *r*_*sens*_, as the sensory signal produced at the object level. Under normal conditions (no TMS), the readout of this sensory signal by DLPFC, *r*_*readout*_, will equal *r*_*sens*_ and will be relayed to aPFC for the confidence judgment. Further, our neural mechanism postulates that the role of aPFC is to integrate the strength of the sensory signal relayed by DLPFC with non-perceptual cues and make the confidence judgement. Within our model, this process can be seen as the addition of metacognitive noise *σ*_*meta*_ at the meta level.

### Modeling the TMS effects

According to our proposed neural mechanism, TMS to DLPFC should influence the magnitude of sensory readout that can be used at the meta level. Our data showed that confidence level decreases following DLPFC TMS, suggesting that the sensory readout decreases in magnitude. We formalized this idea in our computational model as DLPFC TMS leading to a decrease in the magnitude of the sensory readout such that

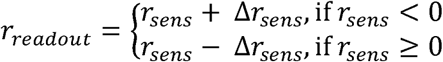

where Δr_sens_ controls the change in the readout. These conditions satisfy the relation |r_*readout*_|*=* |r_sens_|- *Δ*r_sens_ such that the effect of TMS is to reduce the absolute magnitude of *r*_*readout*_ without changing its sign. (As stated above, in cases in which *r*_*sens*_ and *r*_*meta*_ had a different sign - which occurs when |r_sens_| < Δr_sens_ - we constrained the confidence response to equal 1 by setting *r*_*readout*_ = 0.) On the other hand, according to our proposed neural mechanism, TMS to aPFC should affect the level of noise that corrupts the confidence decision. We formalized this idea in our model as aPFC TMS leading to an altered level of metacognitive noise. Since our behavioral results suggested that aPFC TMS increased metacognitive scores only in the second half of blocks, we modeled the effect of aPFC TMS as a decrease in metacognitive noise for the second half of blocks such that *r*_*meta*_ *= N*(r_*readout*_*,(σ*_*meta*_ *-*Δ σ_*meta*_)^*2*^), where *Δa*_*meta*_ controls the change of the metacognitive noise.

To simulate actual data, we set the basic parameters of the model such that μ= 1.74, *σ*_*sens*_ *=1,; σ*_*meta</sub>*_ =0.6*, σ<sub>rneta_base*</sub>=0.15, C__*3*_ =-1.45, C__*2*_ .95, C__*1*_ =-.45 *,c*_*0*_ *=* 0, C_1_ = .45 *,c*_*2*_ *=* 0.95, and c_3_ = 1.45. We set *σ*_*sen*_ = 1 since choosing other values would simply lead to a multiplicative change in all other parameters. The value of |i was chosen based on the average *d′* observed across all subjects. The values for the rest of the parameters were chosen to match the overall performance that we observed in the study. However, the effects of TMS do not depend on the specific numbers and the same qualitative results were observed with a wide range of values of the different parameters. Critically, we used different values of *Δr*_*sens*_ and *Δσ*_*meta*_ for modeling the different TMS conditions. For S1 TMS, we set Ar_sens_ = 0 and Δ<r_meta_ = 0. For modeling DLPFC TMS, we set Δ*r*_*sens*_ *=* 0.072 and *Δσ*_*meta*_ = 0, consistent with the notion that DLPFC TMS should lead to a decrease in the magnitude of the sensory readout for metacognition. Finally, for modeling aPFC TMS, we set Δ*r*_*sens*_ *=* 0 and *Δ σ*_*meta*_*=* 0.65, consistent with the notion that aPFC TMS should change the metacognitive noise.

### Data and code

All data, analysis and simulation files can be downloaded from https://github.com/Medha66/onlineTMS DLPFC aPFC.

## Results

We investigated the specific contributions of DLPFC and aPFC to visual metacognition by e employing an online TMS protocol to disrupt activity within these areas during confidence computation. Subjects indicated the tilt (left/right) of a noisy Gabor patch while simultaneously providing a confidence rating on a four-point scale **(Figure 2A).** On each trial, we delivered a train of three TMS pulses **(Figure 2B)** to DLPFC, aPFC, or S1 (which served as a control site).

**Figure 2:**
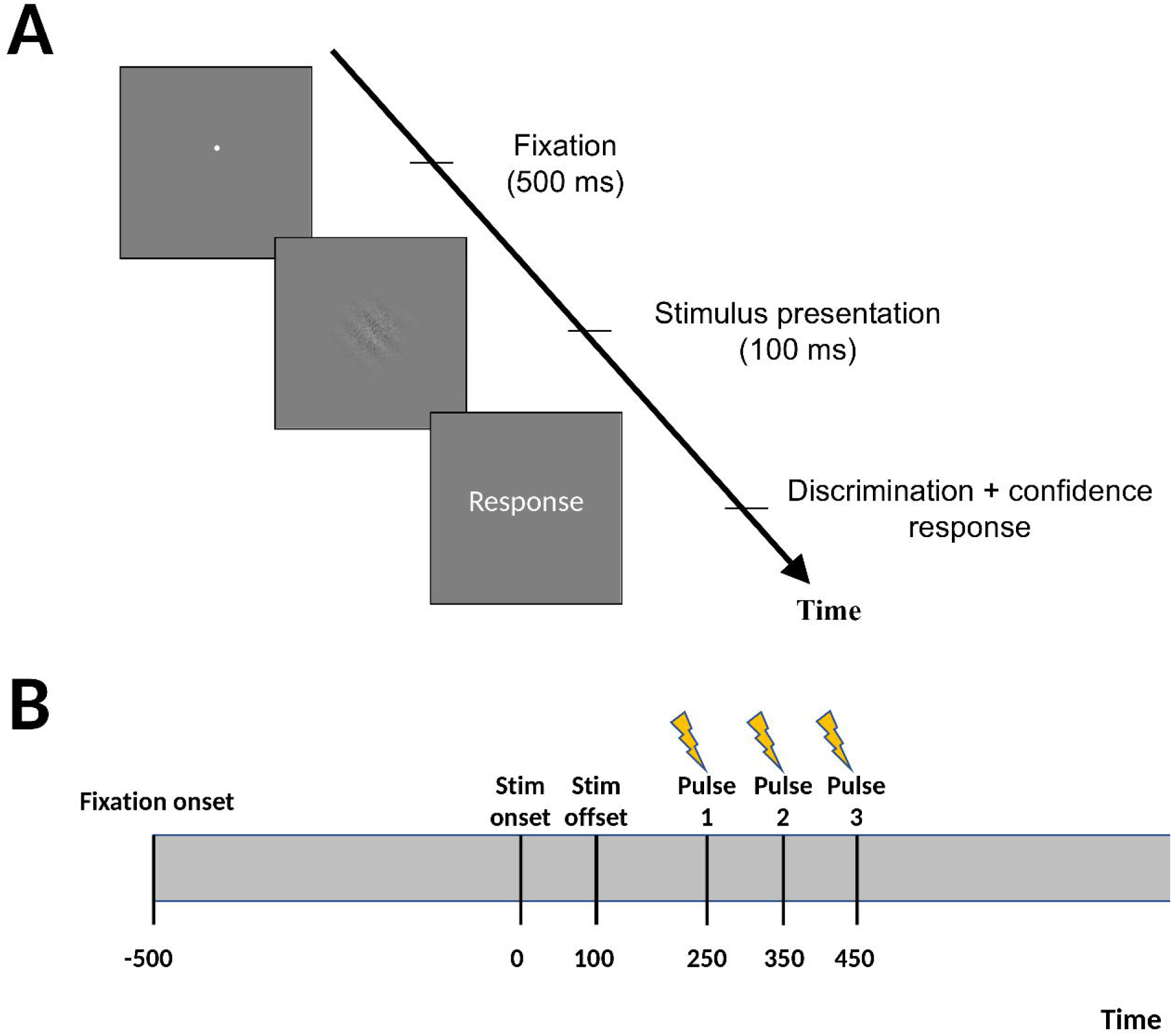
Task. (A) Trial sequence. Each trial began with short fixation (500 ms) followed by the presentation of an oriented Gabor patch (100 ms). Subjects had to simultaneously indicate the tilt (left/right) of the Gabor patch and their confidence on a 1-4 scale. (B) Timeline of TMS delivery. TMS was given as a train of three pulses with inter-pulse interval of 100 ms. The first pulse was delivered 250 ms after onset of the stimulus. Subjects had a mean response time of ∼1000 ms.

As in previous studies on the role of prefrontal cortex in perceptual decision making (Rahnev et al., 2016; Rounis et al., 2010; Ryals et al., 2016), TMS did not influence the overall task performance as measured by accuracy or reaction time (p > 0.05 for all pairwise comparisons between the three sites). These results suggest that the prefrontal cortex is unlikely to be involved in low-level stimulus processing (Rahnev, 2017).

### TMS effect on confidence

Based on our hypothesis regarding the functions of DLPFC and aPFC in confidence generation, we predicted that DLPFC TMS, but not aPFC TMS, would affect subjects′ overall confidence level. The results were consistent with this prediction. Indeed, a one-way repeated measures ANOVA with factor TMS site (S1, DLPFC, and aPFC) demonstrated a significant effect of TMS location on confidence (F(2,17) = 3.68, P = 0.04; **Figure 3).** Pairwise comparisons showed a significant decrease in confidence for DLPFC TMS compared to S1 TMS (difference = 0.09, t(17) = 3.19, P = 0.005). No significant difference was found for comparisons between S1 TMS and aPFC TMS (difference = 0.03, t(17) = 0.83, P = 0.4), implying that overall confidence level was affected only after DLPFC stimulation. The difference in confidence between DLPFC TMS and aPFC TMS was numerically larger than the difference between S1 TMS and aPFC TMS but did not reach significance (difference = 0.06, t(17) = 1.7, P = 0.12).

**Figure 3:**
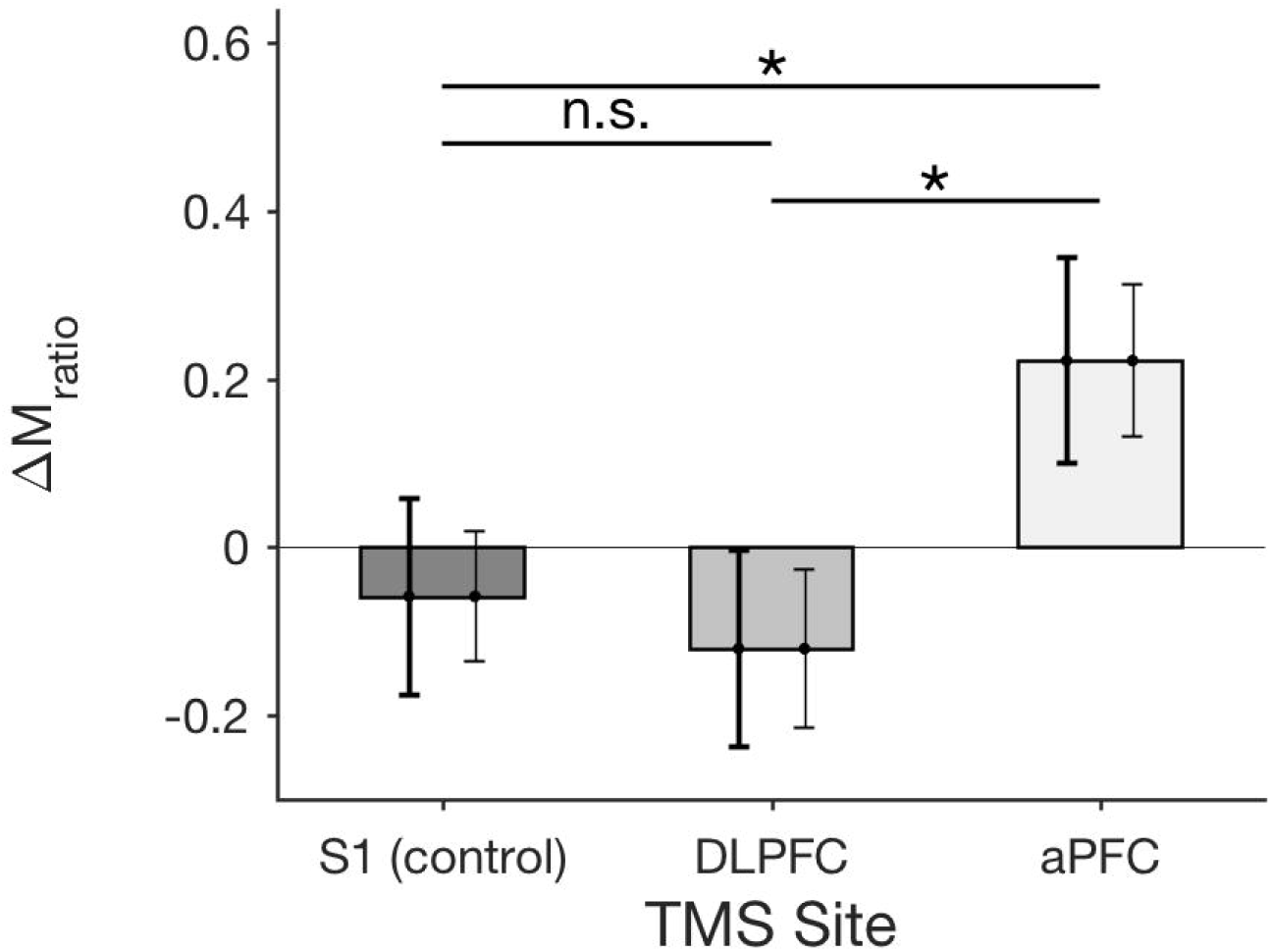
TMS effect on overall confidence level. TMS to DLPFC decreased average confidence while TMS to aPFC did not affect the overall confidence level. The left error bars represent the within-subject standard errors for comparisons with S1 (the error bar for S1 is the same as the one for DLPFC) and are indicative of statistical significance. The right error bars represent the between-subject standard errors and are not indicative of the statistical significance (instead, they show the overall variability in confidence across subjects), n.s. not significant ** P < 0.01.

### TMS effect on metacognitive ability

Based on our hypothesis regarding the functions of DLPFC and aPFC in confidence generation, we predicted that aPFC TMS, but not DLPFC TMS, would affect subjects′ metacognitive ability. To test this prediction, we used *M*_*ratio*_ as a measure of the quality of metacognition (Maniscalco and Lau, 2012). However, a one-way repeated measures ANOVA with factor TMS site (S1, DLPFC, and aPFC) on A/W,_0_ scores showed no main effect of TMS location on metacognitive ability (F(2,17) = 0.3, P = 0.74).

In contrast to these results, previous studies showed that offline TMS to aPFC increased metacognitive scores (Rahnev et al., 2016; Ryals et al., 2016). Therefore, it is possible that the effects of TMS to aPFC become apparent only after a more sustained period of inhibition. To test this possibility, we examined whether metacognitive ability differed between the first and second halves of test blocks. We performed a 2 (time: first vs. second half of test blocks) X 3 (TMS site: S1, DLPFC, and aPFC) repeated measures ANOVA on M_rati0_ values and found a significant interaction between time and TMS site (F(2,1) = 3.9, P = 0.03; **Figure 4).** Further analyses revealed a significant increase in *M*_*ratio*_ for the second half (compared to the first half) of test blocks after aPFC TMS (difference = 0.22, t(17) = −2.44, P = 0.03) but not after S1 TMS (difference = −0.06, t(17) = 0.75, P = 0.47) or DLPFC TMS (difference = −0.12, t(17) = 1.27, P = 0.22). Critically, the difference in *M*_*ratio*_ between the two halves of test blocks was significantly larger for aPFC TMS compared to both S1 TMS (difference = 0.28, t(17) = 2.4, P = 0.028) and DLPFC TMS (difference = 0.34, t(17) = 2.81, P = 0.012). Therefore, TMS increased metacognitive ability for the second half of our blocks and this effect was specific to aPFC.

**Figure 4:**
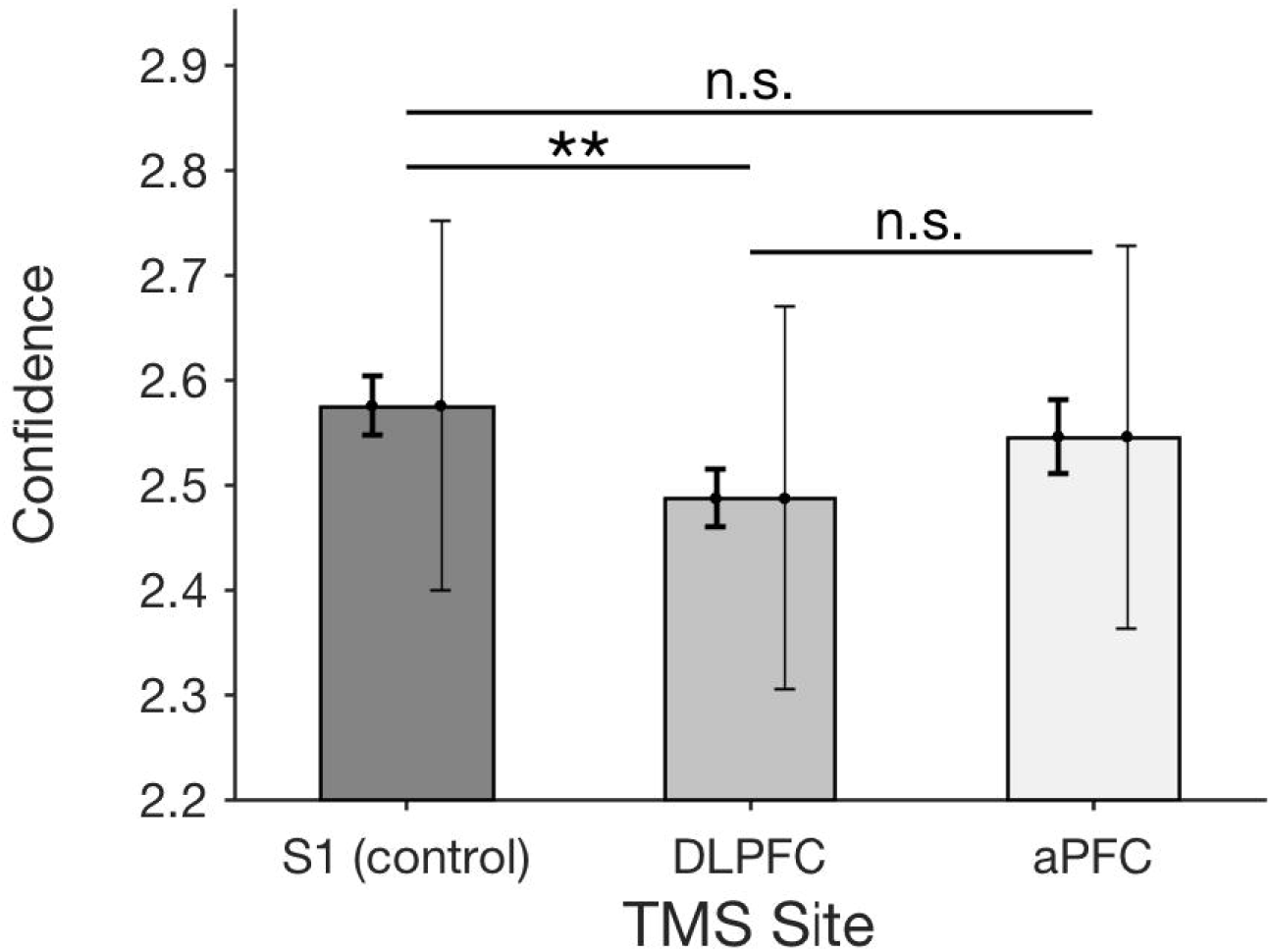
TMS effect on metacognitive ability. TMS to aPFC increased metacognitive ability for the second half compared to the first half of test blocks. No such effect was observed for S1 TMS or DLPFC TMS. Metacognitive ability was operationalized as M_ratio_ (Maniscalco and Lau, 2012). AM_ratio_is the change in M_ratio_from the first half to the second half of a block. The left error bars represent the within-subject standard errors for comparisons with S1 (the error bar for S1 is the same as the one for aPFC) and are indicative of statistical significance. The right error bars represent the within-subject standard errors for comparisons between the first half and second half of blocks and are not indicative of the statistical significance for between-site comparisons, n.s. not significant, * P < 0.05.

We further verified that the changes in *M*_*ra*_*tio* were not driven by changes in the primary task performance *d′.* We performed a two-way repeated measures ANOVA on *d′* with time and TMS location as factors. The results indicated no significant interaction between time and TMS location (F(2,1) = 1.56, P = 0.22). Further, we verified with a paired t-test that the change in *d′* from the first half to the second half of the blocks was not significantly different between aPFC and the control site S1 (t(17) = 1.47, P = 0.16). Although the interaction between time and TMS location did not reach significance for *meta-d′* (F(2,1) = 2.28, P = 0.12), a paired t-test showed that the change in *meta-d′* from the first half to the second half of the blocks was significantly greater for aPFC than S1 (t(17) = 2.27, P = 0.037).

Similarly, we verified that the changes in *M*_*ratio*_ were not driven by changes in confidence. A two-way ANOVA revealed a main effect of time on confidence (F(1,17) = 14.8, P = 0.001) driven by a confidence decrease across all three TMS sites. Critically, the interaction between time and TMS location was non-significant (F(2,1) = 0.42, P = 0.66). A paired t-test showed that the decrease in confidence from first to second half of blocks was not significantly different between aPFC and the control site S1 (t(17) = 0.49, P = 0.63).

These additional results indicate that changes in *d′* and confidence between the two halves of the blocks were similar across the TMS conditions and the observed effect of TMS on *Mratio* for aPFC cannot be explained by these variables.

Simulating the effects of TMS with a hierarchical confidence generation model

The results above confirmed our prediction that disrupting DLPFC would affect average confidence, while disrupting aPFC would affect metacognitive ability. This prediction was based on the hypothesis that DLPFC reads out the strength of the sensory signal and relays it to aPFC, which translates it into a confidence judgment by also incorporating non-perceptual factors. To test whether these mechanistic effects can indeed reproduce our results, we implemented them in simulations of a computational model of confidence generation.

The model that we developed is based on the common assumption of the existence of independent sensory and metacognitive noise (Mueller and Weideman, 2008; De Martino et al., 2013; Rahnev et al., 2016; Bang et al., 2017; van den Berg et al., 2017). The two noise stages lead to separate representations for object- and meta-level judgments **(Figure 5A).** At the object level, the stimulus is corrupted by sensory noise and the resulting signal is used to make a perceptual decision. To make the confidence judgment, the signal strength from the object level is read out at the meta level. The final confidence decision is based on the sensory readout, as well as other factors such as the history of confidence responses (Rahnev et al., 2015), perceived attentional state, etc. We modeled all of these influences collectively as the addition of metacognitive noise.

**Figure 5.**
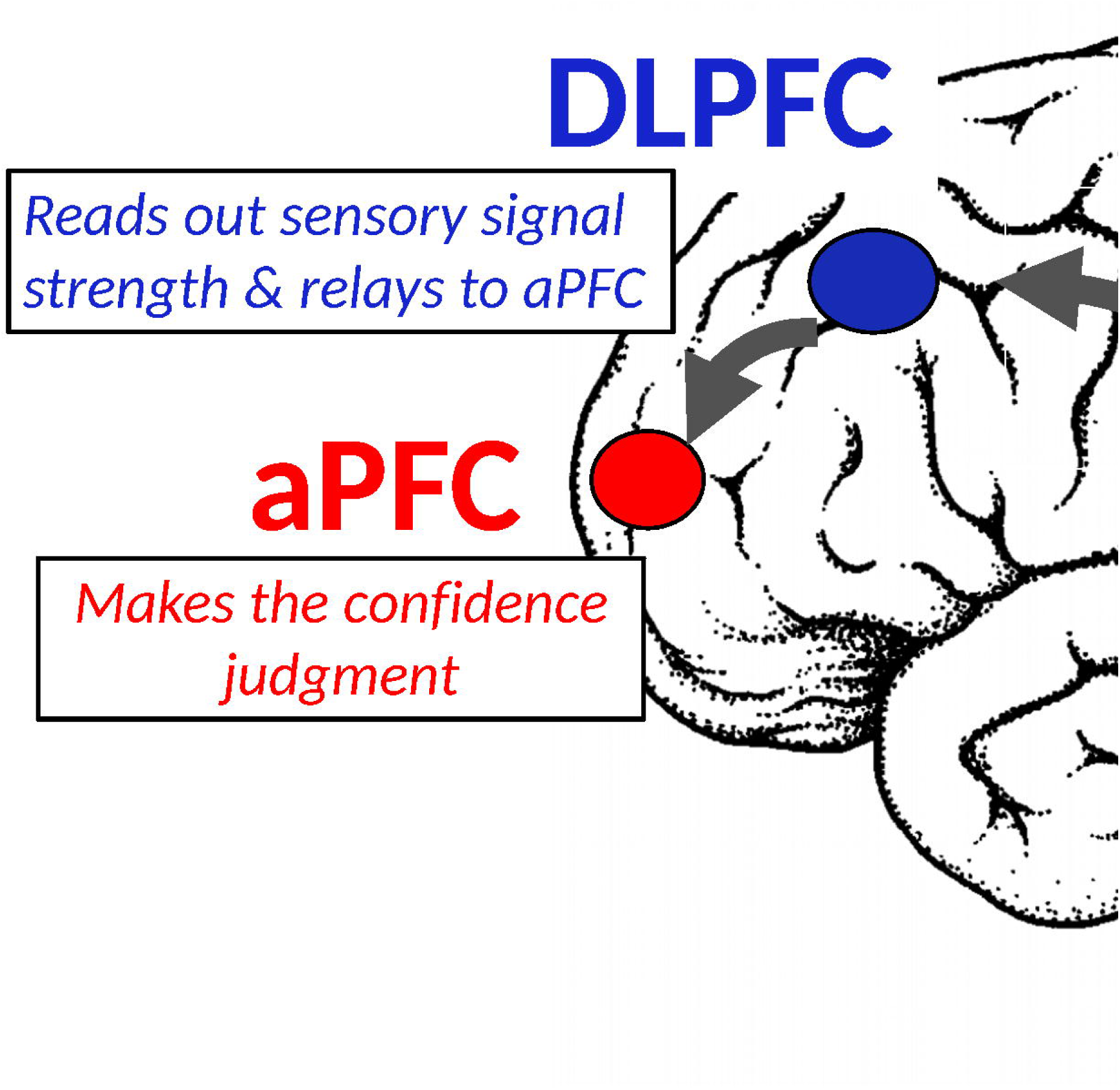
A computational model of confidence generation. (A) The sensory signal (r_sens_) available at the decision level is read out (r_readout_) to the metacognitive level and additional noise (σ_meta_) is added before obtaining the confidence signal (r_meta_). The perceptual decision is based on the sensory signal r_sens_, while the confidence judgment is based on the confidence signal r_meta_. Consistent with the hypothesized roles of DLPFC and aPFC in confidence computation, we modelled the effect of DLPFC TMS as a signal loss from the readout (quantified as Ar_sens_; boxed equation on the left), and the effect of aPFC TMS as lowered metacognitive noise (quantified as Ao_meta_; boxed equation on the right). (B-C) Model simulations show that decreasing the magnitude of the readout decreases the overall confidence level (panel B) but does not influence metacognitive ability (panel C). Conversely, decreasing the amount of metacognitive noise in the second half of test blocks has a small effect on average confidence (panel B) but a large effect on increasing the difference in metacognitive ability between the first and second half of blocks (panel C). These results mirror the effects of TMS to DLPFC and aPFC in our data (see Figures 3 and 4).

Within this architecture, our proposed effects of inhibiting DLPFC and aPFC can be operationalized as DLPFC TMS affecting the strength of the sensory readout, and aPFC TMS affecting the level of metacognitive noise **(Figure 5A;** boxed equations). Quantitatively, we modeled the effect of TMS to DLPFC as a loss of the sensory readout at the meta level and the effect of TMS on aPFC as a decrease in metacognitive noise (see Methods).

Simulations of our computational model faithfully reproduced the TMS effects for both overall confidence level **(Figure 5B)** and metacognitive ability **(Figure 5C).** Therefore, within this established architecture of hierarchical confidence generation, our TMS results can be recreated by assuming a role for DLPFC in the reading out the sensory signal strength at the meta level, and a role for aPFC in making the final confidence judgment based on a combination of perceptual and non-perceptual factors.

## Discussion

We sought to determine the distinct roles of subregions of the prefrontal cortex in visual metacognition. Previous research identified the dorsolateral and anterior prefrontal cortex (DLPFC and aPFC) as critical to metacognitive computations but a mechanistic understanding of their functions in confidence judgments is still lacking (Shimamura, 2000a; Fleming and Dolan, 2012). We proposed a neural mechanism for confidence computation where DLPFC reads out the sensory signal strength and relays it to aPFC, while aPFC makes the confidence judgment by integrating this readout with non-perceptual factors. Based on this architecture, we predicted that disrupting DLPFC would affect average confidence (without affecting metacognitive ability), while disrupting aPFC would affect metacognitive ability (without affecting confidence). A causal intervention with online TMS confirmed these predictions. Further, we simulated a confidence generation model that incorporated our hypothesized neural mechanism and successfully reproduced the observed behavioral results. These findings establish the existence of independent causal contributions of DLPFC and aPFC to confidence generation and suggest specific mechanistic roles for these prefrontal sites. Further, they suggest that a significant portion of confidence computation in PFC takes place 250-450 ms following stimulus onset.

### Role of DLPFC in confidence computation

Our experiment tested the hypothesis that the role of DLPFC in confidence computation is to read out the strength of the sensory signal and relay it to aPFC. We derived this hypothesis from previous studies, which found that DLPFC activity is related to the level of confidence but not to metacognitive ability (Fleck et al., 2005; Henson et al., 2000; Lau & Passingham, 2006). This proposed function of DLPFC in reading out the sensory signal strength is consistent with the view that DLPFC maintains, reroutes, and facilitates manipulations of sensory information (Shimamura, 2000b; Fleming and Dolan, 2012).

The correlation between DLPFC activity and confidence level has received different interpretations. Henson et al. (2000) hypothesized that DLPFC activity reflects retrieval monitoring in a memory task. Fleck et al. (2005) suggested a general role for DLPFC in information monitoring during decision making. Finally, Lau & Passingham (2006) theorized that DLPFC plays a role in conscious perception. Our proposal - that the role of DLPFC in confidence computations is to read out the strength of the sensory signal - is not necessarily at odds with these previous theories. Instead, here we specify a precise computational role for DLPFC in the domain of confidence generation.

There has been some controversy about whether DLPFC is involved more directly in confidence computation. Rounis et al. (2010) delivered bilateral TBS to DLPFC and reported a decrease in mean visibility as well as metacognitive performance. These findings have been controversial with Bor et al. (2017) arguing that they could not replicate them, while Ruby et al. (2017) disputing Bor et al.′s exclusion criteria and arguing that the original effects replicate under different exclusion criteria. While our study certainly has implications about the role of DLPFC in metacognition, it is not clear whether it can be used to inform the above debate. Indeed, both studies above (Bor et al., 2017; Rounis et al., 2010) targeted a relatively posterior portion of DLPFC, while we targeted a relatively anterior DLPFC region. DLPFC is anatomically large and it is likely that its different sub-regions have different functions. Other important differences between ours and the two studies above include the use of an online vs. offline TMS protocol, unilateral vs. bilateral stimulation, and confidence vs. visibility ratings with each of these factors making direct comparisons difficult.

### Role of aPFC in confidence computation

Our experiment tested the hypothesis that the role of aPFC in confidence computation is to decide the exact value of the confidence rating based on both the sensory readout relayed by DLPFC and other, non-perceptual factors. In line with this hypothesis, many previous studies have found a link between aPFC and metacognitive ability (Fleming et al., 2010, 2012b, 2014; Yokoyama et al., 2010; McCurdy et al., 2013; Rahnev et al., 2016; Ryals et al., 2016; Allen et al., 2017). Our proposal that aPFC is the seat of metacognitive computation is also consistent with the view that aPFC is at the highest level in the cognitive and perceptual decision-making hierarchy (Badre and D′Esposito, 2009; Fleming and Dolan, 2012; Rahnev,2017).

A wide range of higher-order functions in the domains of memory, cognition, and perceptual decision making have been attributed to aPFC. These functions include top-down manipulations of working memory representations, switching between task sets, attentional allocation to sub-goals, and relational integration (Koechlin et al., 1999; Kaas et al., 2007; Domenech and Koechlin, 2015; Lara and Wallis, 2015; Parkin et al., 2015). Ramnani & Owen (2004) integrate these theories into a common framework which proposes that aPFC recruitment facilitates the coordination of information processing from separate mental processes towards a higher goal. This view is fully consistent with our theory′s implication of aPFC in generating metacognitive computations. Indeed, assessing the confidence in one′s own perceptual decisions requires the integration of both perceptual and non-perceptual factors (Fleming & Dolan, 2012).

We found that TMS influenced metacognitive ability only for the second half of trials within a block. It appears that a sustained period of inhibition may be required in order to influence metacognitive ability. Indeed, previous studies that successfully manipulated metacognitive ability (Rahnev et al., 2016; Ryals et al., 2016) employed offline TMS, which involves a sustained period of stimulation. More research is needed to determine whether TMS may interact differently with the unique cytoarchitectonic characteristics of aPFC (Semendeferi et al., 2001).

Disrupting the activity of aPFC during confidence computation improved metacognitive performance. While such an improvement appears surprising at first, it is consistent with previous studies that found increases in metacognitive ability after offline TMS to aPFC (Rahnev et al., 2016; Ryals et al., 2016). One possible explanation for this increased ability is that aPFC TMS increased the attentional resources for the confidence decision. However, increased attentional resources could be expected to also lead to increases in *d′* and confidence but aPFC TMS had no effect on either of these measures. Another possibility is that TMS might have inhibited the influence of certain factors that are detrimental to metacognition. For example, people consider their confidence history while making a confidence judgement, a phenomenon called confidence leak (Rahnev et al., 2015). Confidence ratings may also be contaminated by other factors such as arousal (Allen et al., 2016), action fluency (Fleming et al., 2015), etc. The use of these extra factors generally decreases metacognitive ability in laboratory settings (Rahnev et al., 2015). Therefore, the improvement of metacognitive ability with aPFC TMS in our study may stem from the reduced use of some of these non-perceptual factors in confidence generation.

### Computational model

We built a computational model that instantiates the hypothesized neural mechanism regarding the roles of DLPFC and aPFC. It is important to note that while the TMS data provide support for the proposed neural mechanism, our experiment was not designed to corroborate the computational model directly. Instead, the role of the computational model was to verify that the substantive claims made by our neural mechanism could indeed lead to the pattern of behavioral results that we observed. We have explored the plausibility of our computational model elsewhere (Bang et al., 2017).

We modeled the effect of TMS on aPFC and DLPFC as a decrease in metacognitive noise and a decrease of signal in the sensory readout (*r*_readout_), respectively. The modeling choice for aPFC TMS is natural given that, within our model, metacognitive ability is controlled by the metacognitive noise parameter. However, the effects of DLPFC TMS on decreased confidence can also be explained as a shift in the confidence criteria. The reason we do not favor this explanation is because it is unclear why TMS would shift the criteria in one direction and not the other. Specifically, we are not aware of any mechanism that predicts that TMS would increase the confidence criteria (in order to decrease confidence). Instead, our explanation - that TMS causes a loss of signal, which leads to a confidence decrease - relates more naturally to the expected effect of TMS, which is to disrupt neural activity.

### Conclusion

Our results show that TMS produced distinct effects on confidence measures depending on which prefrontal site was stimulated: TMS to DLPFC decreased confidence, while TMS to aPFC increased metacognitive ability for the second half of the experimental blocks. This dissociation confirms our hypothesis that DLPFC and aPFC have distinct roles in visual metacognition. Further, it supports our hypothesized neural mechanism, according to which DLPFC reads out the sensory signal strength and relays it to aPFC for the confidence computation. Simulations of a confidence generation model based on our neural mechanism reproduced the observed TMS effects and thus corroborated this mechanism. Together, our results uncover the functional organization of PFC for confidence computations.

## Acknowledgments

We thank Ji Won Bang for help with the experiment setup and Lindley Hudson for assistance with experiment preparation and subject recruitment. This work was funded by a startup grant to D.R. from the Georgia Institute of Technology.

